# Isolation and characterization of electrochemically active subsurface Delftia and Azonexus species

**DOI:** 10.1101/046979

**Authors:** Yamini Jangir, Sarah French, Lily M. Momper, Duane P. Moser, Jan P. Amend, Mohamed Y. El-Naggar

**Author notes:** Correspondence: Mohamed Y. El-Naggar, University of Southern California, 920 Bloom Walk, Los Angeles, CA 90089-0484, USA.

## Abstract

Continental subsurface environments can present significant energetic challenges to the resident microorganisms. While these environments are geologically diverse, potentially allowing energy harvesting by microorganisms that catalyze redox reactions, many of the abundant electron donors and acceptors are insoluble and therefore not directly bioavailable. Extracellular electron transfer (EET) is a metabolic strategy that microorganisms can deploy to meet the challenges of interacting with redox-active surfaces. Though mechanistically characterized in a few metal-reducing bacteria, the role, extent, and diversity of EET in subsurface ecosystems remains unclear. Since this process can be mimicked on electrode surfaces, it opens the door to electrochemical techniques to enrich for and quantify the activities of environmental microorganisms *in situ*. Here, we report the electrochemical enrichment of microorganisms from a deep fractured-rock aquifer in Death Valley, California, USA. In experiments performed in mesocosms containing a synthetic medium based on aquifer chemistry, four working electrodes were poised at different redox potentials (272, 373, 472, 572 mV vs. SHE) to serve as electron acceptors, resulting in anodic currents coupled to the oxidation of acetate during enrichment. The anodes were dominated by *Betaproteobacteria* from the families *Comamonadaceae* and *Rhodocyclaceae.* A representative of each dominant family was subsequently isolated from electrode-associated biomass. The EET abilities of the isolated *Delftia* strain (designated WE1–13) and *Azonexus* strain (designated WE2–4) were confirmed in electrochemical reactors using working electrodes poised at 522 mV vs. SHE. The rise in anodic current upon inoculation was correlated with a modest increase in total protein content. Both genera have been previously observed in mixed communities of microbial fuel cell enrichments, but this is the first direct measurement of their electrochemical activity. While alternate metabolisms (e.g. nitrate reduction) by these organisms were previously known, our observations suggest that additional ‘hidden’ interactions with external electron acceptors are also possible. Electrochemical approaches are well positioned to dissect such extracellular interactions that may be prevalent in the subsurface.

## INTRODUCTION

The observation that microorganisms permeate subsurface environments down to kilometer depths highlights the astounding range of metabolic strategies that our planet’s unseen majority must wield to survive in such environments (Edwards et al., 2012). Far removed from the resources available to surface life, subsurface microorganisms must contend with heterogeneous habitats that may present a shortage of organic carbon and other nutrients necessary for biomass, cope with environmental stressors (extremes of temperature, pressure, pH, and radioactivity), as well as extract energy for metabolic activity despite the scarcity of soluble electron donors and acceptors (Parnell and McMahon, 2016). However, while much emphasis is usually placed on soluble redox couples because they are directly bioavailable for intracellular reactions, it is important to note that the subsurface geological environment also offers insoluble electron donors and acceptors in the form of redox active elements (e.g. S, Fe, Mn) in minerals associated with sediments and rocks (Nealson et al., 2002; Bach and Edwards, 2003; Edwards et al., 2005; Fredrickson and Zachara, 2008; Orcutt et al., 2011; Southam, 2012). Microbes capable of performing extracellular electron transfer (EET), allowing cellular electron flow from or to inorganic minerals, may therefore have a selective advantage for energy acquisition by uptake or disposal of electrons at the biotic-abiotic interfaces prevalent in subsurface habitats.

To date, however, the mechanistic bases of EET are well characterized in only two model bacterial systems, the*Deltaproteobacterium, Geobacter* and the *Gammaproteobacterium, Shewanella* (Lovley and Phillips, 1988; Myers and Nealson, 1988). The elucidation of detailed EET mechanisms has benefited from electrochemical approaches where electrodes substitute for the insoluble minerals that these organisms can utilize as terminal electron acceptors in sedimentary environments (Bond, 2010). The identified mechanisms range from the use of multiheme cytochrome ‘conduits’ that mediate electron transfer through the cellular periplasm and outer membrane to external surfaces (Myers and Myers, 1992; Hartshorne et al., 2009; White et al., 2013), electron transfer to more distant surfaces along bacterial nanowires (Reguera et al., 2005; Gorby et al., 2006), and possibly soluble redox shuttles proposed to link the cellular redox machinery to external surfaces by diffusion without the need for direct cell-surface contact (Marsili et al., 2008; von Canstein et al., 2008).

Beyond the well characterized model systems, however, electrochemical enrichments coupled with 16S rDNA-based surveys from a variety of environments, especially marine sediments, suggest that more physiologically and phylogenetically diverse microorganisms are capable of using electrodes as electron acceptors (Holmes et al., 2004; Rabaey and Verstraete, 2005; Reimers et al., 2006; White et al., 2009; Bond, 2010). It is important to note that the survival or enrichment of a microbe on an electrode is not itself evidence of EET ability, since operational conditions, including potential oxygen leakage or the use of fermentable substrates, may support subpopulations of heterotrophic aerobes or fermentative organisms, for instance. This makes isolation and electrochemical characterization fundamental to investigating EET processes. As such, several enrichments have led to the isolation of pure cultures that do interact with electrodes (Wrighton et al., 2008; Xing et al., 2008; Zuo et al., 2008; Fedorovich et al., 2009), indicating that microbial electrochemical activity at redox-active surfaces may be advantageous in a wide variety of habitats. Indeed, the ability to transfer electrons to anodes has now been confirmed in multiple bacterial phyla (Fedorovich et al., 2009). Recent studies even point to a great diversity of microbes that can perform electron *uptake* from electrodes, including acetogenic bacteria (Nevin et al., 2011), methanogenic archaea (Deutzmann et al., 2015), as well as iron-and sulfur-oxidizing bacteria (Summers et al., 2013; Bose et al., 2014; Rowe et al., 2015). It now appears that microbial EET may be widespread in nature, and that electrode-based techniques are critical for both shedding light on the phylogenetic diversity and for identification and mechanistic measurements of organisms whose electrochemical activity would have been missed using traditional cultivation techniques.

Our study was motivated by both the apparent extent of EET in nature, and the particular relevance to the continental subsurface that may present opportunities for microbial interaction with redox-active abiotic surfaces. Here, we utilized electrodes poised at anodic (electron-acceptor) redox potentials for enrichment of electrochemically-active bacteria from a deeply sourced artesian well that is supplied by a large regional flow system in Death Valley, California, USA. The enrichments at different anodic potentials, subsequent isolation of pure cultures, and small currents observed by electrochemical testing of these pure cultures under well-defined conditions, suggest that isolated *Delftia* and *Azonexus* strains may gain an advantage by passing electrons to external surfaces. Our results contribute to the emerging view that microbial electrochemical activity is a widespread phenomenon that may impact survival and activity in energy-limited conditions.

## MATERIALS AND METHODS

### SAMPLING SITE AND INITIAL ENRICHMENT

The sampling site (Nevares Deep Well 2-NDW2) was drilled without drilling fluid in 2009, penetrating the Nevares Spring Mound, a prominent travertine assemblage located at the western foot of the Funeral Mountains in Death Valley, California, USA (Moser, 2011). The hole was drilled to 103 meters below land surface, but cased only to a depth of 18 m, below which it is open and intersects with fractures of the Furnace Creek Fault Zone. Once in the fault, the hole opens into a rubble zone from which artesian flow (~300 L/min) emanates (Michael King, Hydrodynamics Group LLC, and Richard Friese, National Park Service - personal communication). After flushing the hole with 200 well volumes, freshwater samples were collected for aqueous chemistry (Table 1)) and microbiological analyses. Aqueous geochemical analyses (Table 1) were performed at the Analytical Lab San Bernardino (ALSB). Briefly, anions were measured according to Environmental Protection Agency method 300.0 using ion chromatography. Cations and metals were determined using Inductively Coupled Plasma Mass Spectrometry (ICP-MS), and Atomic Absorption (AA), respectively. Water samples were used for an initial enrichment in a Down-flow Hanging Sponge (DHS) reactor (Imachi et al., 2011), targeting Fe-and Mn-reducing microbes. The minerals 2–line ferrihydrite (containing Fe-III) and 3–MnO_2_ (containing Mn-IV) were freshly made for inoculation of the DHS reactor (Myers and Nealson, 1988; Schwertmann and Cornell, 2000). Aqueous medium was designed to mimic the geochemical composition *in situ* (Table 1). Specifically, 1 L of medium contained: 0.1g NH_4_—Cl, 23 0.1g NH_4_SO_4_, 0.2g KH_2_PO_4_, 0.1g MgSO_4_ and 0.2g CaCl_2_. After autoclaving, pH was set at 7.2 with 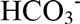buffer and DSMZ medium 141 trace metals and vitamins were filter sterilized into medium. The gas phase was composed of 80:20 v:v H_2_:CO_2_. Aqueous and gaseous media were provided at 0.1 mL per minute via a peristaltic pump (Cole-Parmer Masterflex L/S, model number 7551-10) and media was kept anaerobic with 100 µM Ti-NTA (freshly made from 20% TiCl_3_ and nitrilotriacetic acid (NTA) in saturated Na_2_CO_3_). Resazurin (0.001%) was used as a redox indicator.

**Table 1.** Chemical composition of water collected from Nevares Deep Well 2 (Death Valley, 1 CA, USA).

|  | NDW2 |
| --- | --- |
| Depth (m) | 103 |
| pH | 7 |
| Temperature (°C) | 31 |
| Conductance (μS/cm) | 790 |
| Dissolved Constituents |  |
| dO<sub>2</sub> (mg/L) | 0.4 |
| SO<sub>4</sub><sup>2-</sup> (mg/L) | 170 |
| Na<sup>+</sup> (mg/L) | 160 |
| Ca<sup>2+</sup> (mg/L) | 44 |
| Mg<sup>2+</sup> (mg/L) | 21 |
| DOC (mg/L) | 0.12 |
| Fe<sup>2+</sup> (mM) | 0.01 |
| H<sub>2</sub>S (mg/L) | 0.0 |
| Cl<sup>-</sup> (mg/L) | 35.8 |
| Br<sup>-</sup> (mg/L) | 0.25 |
| NO<sub>3</sub><sup>-</sup> (mg/L) | 0.13 |
| PO<sub>4</sub><sup>3-</sup> (mg/L) | 0.06 |
| NO<sub>2</sub><sup>-</sup> (mg/L) | < 0.5 |

### ELECTROCHEMICAL ENRICHMENT BIOREACTOR

The electrochemical enrichment bioreactor (see schematic in Figure 1) was constructed from a polypropylene wide-mouth jar (1 L), which contained a PTFE assembly of four threaded rods, each of which supported a working electrode (WE) composed of 3 x 2 cm carbon cloth (PW06, Zoltek, St. Louis, MO, USA). The bioreactor was a standard half-cell that contained, in addition to the four carbon cloth working electrodes, a common platinum counter electrode (VWR, PA, USA) and a common Ag/AgCl reference electrode (1M KCl, CH Instruments, TX, USA). During enrichment, the four working electrodes (WE1, WE2, WE3, and WE4) were poised at 272, 372, 472, 572 mV vs. SHE, respectively, using a four-channel potentiostat (EA164 Quadstat, EDaq, USA). All electrical connections were made using insulated titanium wires. The complete set-up was autoclaved with working and counter electrodes, and the ethanol-sterilized reference electrode was inserted before use. The bioreactor was fed using a media reservoir. Both the bioreactor and media reservoir were continuously purged with sterile, filtered N_2_ gas to maintain anaerobic conditions.

**Figure 1.**
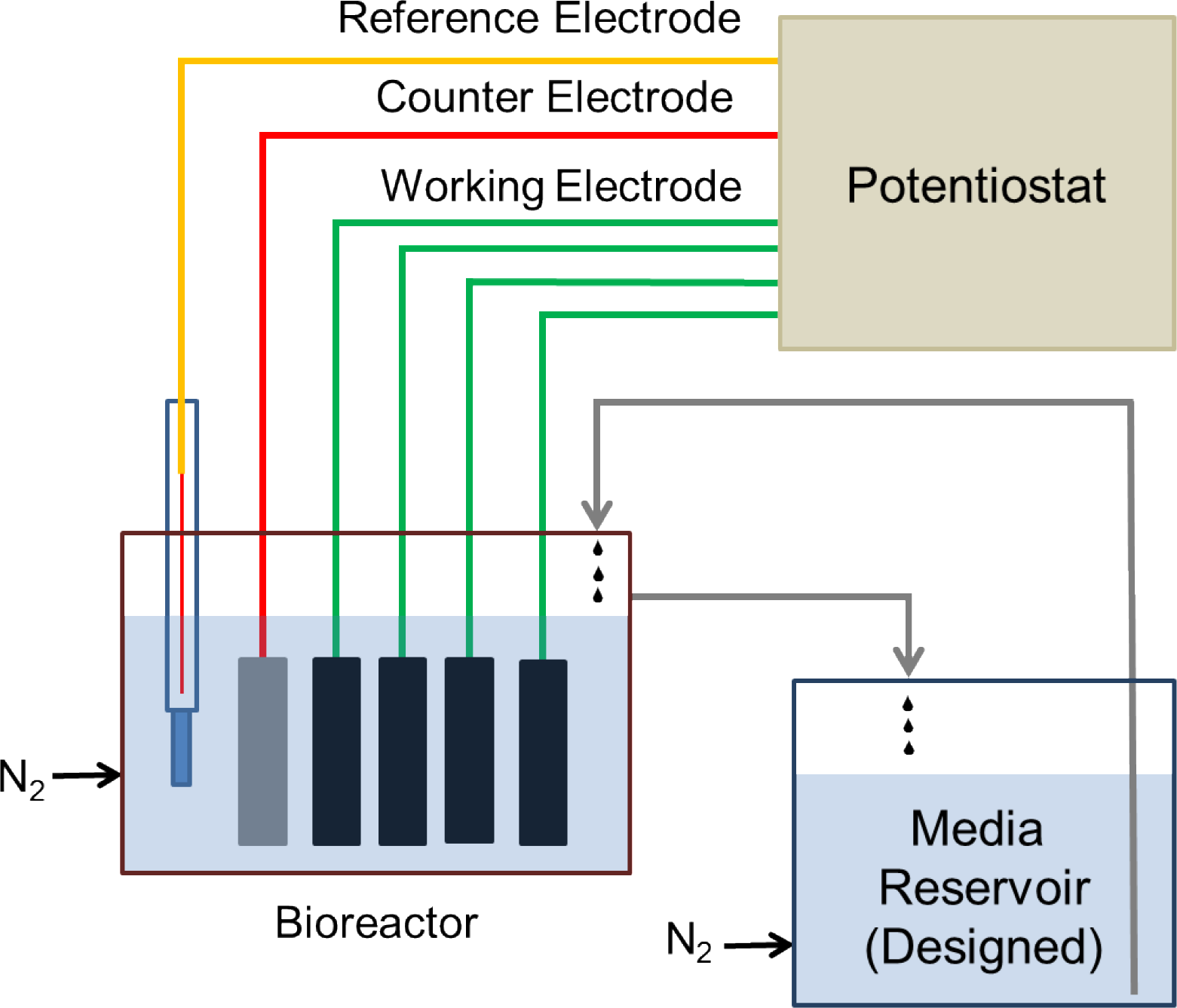
**Schematic diagram of the electrochemical enrichment reactor** The four carbon cloth working electrodes were poised at 272 mV, 372 mV, 472 mV, 572 mV vs. SHE, respectively, using a 4-channel potentiostat. The reactor and medium reservoir were continuously purged with filter-sterilized N2 gas to maintain anaerobic conditions.

During the first phase of electrochemical enrichment, designed to match the conditions of the DHS reactor, DHS effluent was used as an inoculum and the enrichment medium consisted of autoclaved NDW2 well water supplemented with (L^-1^): NH_4_Cl, 0.025 g; (NH_4_)_2_SO_4_, 0.132 g; KH_2_PO_4_, 0.095 g; with DSMZ medium 141 trace metals and vitamins. The media reservoir was constantly bubbled with sterile filtered N2 to maintain anaerobic conditions, while the bioreactor was bubbled with 80:20 v:v H_2_:CO_2_ to maintain similar conditions as the DHS bioreactor. This enrichment was performed for 30 days, and WE4 (+572 mV vs. SHE) was used as an inoculum for a batch enrichment of Fe(III)-reducers in the same medium using 10 mM sodium acetate as electron donor and 5 mM Fe(III)-NTA as electron acceptor. After growth (increase in cell counts confirmed by DAPI staining) and reduction of Fe(III)-NTA (detected visually by color change of Fe(III) to colorless Fe(II), and confirmed by ferrozine assay) (Stookey, 1970), the batch culture was further enriched in the electrochemical bioreactor. During this second electrochemical phase, NDW2 medium was designed with sodium acetate (10 mM) serving as electron donor, and basal salts and nutrients (L^-1^): MgSO_4_.2H_2_O, 0.19 g; CaCl_2_.H_2_O, 0.15 g; NH_4_Cl, 0.025 g;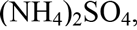,0.132 g; KH_2_PO_4_, 0.095 g; vitamins and trace minerals (Kieft et al., 1999). Both the media reservoir and the bioreactor were bubbled constantly with N_2_ to maintain anaerobic conditions. This enrichment was operated continuously for 6 months, with a media reservoir change performed every 10-30 days. The dilution rate in the bioreactor amounted to one reactor volume per day. Acetate concentration was monitored by high performance liquid chromatography (HPLC), as described previously (Bretschger et al., 2010).

### BACTERIAL COMMUNITY ANALYSIS OF ENRICHMENTS

Working electrode sections (1 x 1 cm) and planktonic cells (500 µL) from the electrochemical bioreactor were suspended with 700 µL lysozyme buffer (25 mM Tris HCl, pH 8.0, 2.5 mM EDTA with 2 mg/mL lysozyme) in a microcentrifuge tube, followed by heating on a microtube shaking incubator at 65°C for 10 min. The mixture was vortexed at maximum speed for 10 min and incubated at 65°C with vigorous shaking for 20-30 min after adding 700 µLTESC buffer (100 mM Tris HCl pH 8.0, 100 mM EDTA pH 8.0, 1.5 M NaCl, 1% w/v cTAB, 1% w/v SDS, adjusted final pH of TESC buffer to 10, added 100 µg/mL Proteinase K just before use) for cell lysis. Further, vortexing for 10 s was performed, followed by incubation on ice for 10-20 min. Cellular debris was removed by centrifugation at ~18000 x g for 10 min. The supernatant was mixed with 25:24:1 phenol:chloroform:isoamyl alcohol. The mixture was spun down at 14000 rpm for 10 min and the supernatant was transferred to a fresh microcentrifuge tube. This collected supernatant was mixed with 750 p,L of isopropanol (no vortexing) and incubated at room temperature for 5 min. Final spin was performed at ~18000 x g for 10 min and the pellet was washed with 95% ice-cold ethanol. The above step of centrifuging the pellet followed by washing was repeated 2-3 times. The pellet was dried overnight until no liquid was left. The extracted DNA was suspended in 100 µL of TE buffer (pH 8.0, 10 mM Tris-HCl and 1 mM EDTA), and DNA concentration and purity were assessed using a nanodrop spectrophotometer.

The near-complete bacterial 16S rRNA gene was amplified using the extracted DNA with primers 8f: 5’ - AGA GTT TGA TCC TGG CTC AG -3’ and 1492r: 5’-GGT TAC CTT GTT ACG ACT T-3’ to perform microbial community analysis (Turner et al., 1999). PCR amplification was carried out in 20 µL reaction volumes using TaKaRa ExTaq DNA Polymerase (TaKaRa Biosciences, Mountainview, CA), and products cloned using pCR^TM^2.1-TOPO (Invitrogen, Carlsbad, CA). Transformed Giga competent cells *(E. coli* DH5a) were grown overnight in Luria-Bertani (LB) agar plates containing Isopropyl β-D-1-thiogalactopyranoside (IPTG), 5-bromo-4-chloro-3-indolyl-β-galactopyranoside (X-gal), and 50 εg/mL kanamycin. Approximately 40 recombinant colonies (white) were picked from each sample (originating from the WE1, WE2, WE3, WE4, and planktonic), and grown in LB broth with kanamycin overnight. Plasmid extraction was performed using the PureLink quick plasmid DNA miniprep kit (Invitrogen, Carlsbad, CA). This DNA was sent for unidirectional sequencing to Genewiz (South Plainfield, NJ). The sequences (600 bp) were analyzed using Geneious software and compared to other published sequences using Naïve Bayesian Classifier provided by the Ribosomal Database Project (RDP) classifier tool.

High-throughput pyrosequencing was also applied to the same DNA extracts used to construct clone libraries for bacterial community analysis across the samples. Variable region V4 of the bacterial 16S rRNA gene amplicon was amplified using barcoded fusion primer 515f: 5’-GTG CCA GCM GCC GCG GTA A-3’ and 806r: 5’-GGA CTA CHV GGG TWT CTA AT-3’. All samples were pooled in equimolar concentrations and purified using Agencourt Ampure beads (Angencourt Biosciences Corporation, MA, USA). Samples were sequenced at MR DNA (Shallowater, TX, USA) with Roche 454 FLX titanium instrument. The resulting data were processed using MOTHUR (Schloss et al., 2009). Sequences were depleted of barcodes and the primer, followed by trimming to remove sequences with any ambiguous base calls and homopolymer runs greater than 8 bp. Average sequence lengths of 260 bp and a total of approximately 12,000 unique sequences were obtained for all samples. The trimmed sequences were aligned with RDP infernal aligner (Nawrocki et al., 2009), chimeras were removed using UCHIME (Edgar et al., 2011), and a distance matrix was created. The sequences were clustered to identify unique operational taxonomical units (OTUs) at the 97% level, and taxonomy was assigned using the RDP classifier. The raw sequences have been uploaded to NCBI SRA database (accession number: SRP071268).

### ISOLATION AND ELECTROCHEMICAL MEASUREMENTS OF PURE CULTURES

Electrode-attached biomass from each WE was streaked on R2A agar plates (Reasoner and Geldreich, 1985) to obtain colonies at 30°C. Morphologically-distinct colonies were restreaked on fresh R2A agar plates, resulting in multiple isolates. For taxonomic classification, isolates were grown to late exponential phase in liquid R2A at 30°C and DNA extraction performed using the UltraClean^®^ Microbial DNA Isolation kit (Mo Bio laboratories, Carlsbad, CA). The bacterial 16S rRNA gene was PCR-amplified from extracted DNA with primers 8f and 1492r, and the purified PCR product (PureLink^®^ PCR Purification Kit, Life Technologies, CA, USA) was sequenced (Genewiz, South Plainfield, NJ, USA) from the 1492r primer. The 600 bp length sequences of the isolated strains have been deposited to Genbank (accession numbers: KU836931, KU836932).

Chronoamperometry measurements for each isolate were performed in standard three-electrode glass electrochemical cells (50 mL volume). Carbon cloth (1 x 1 cm) was used as the working electrode (WE), with platinum wire as counter electrode, and a 1M KCl Ag/AgCl reference electrode. Each isolate was grown to stationary phase aerobically from frozen stocks (−80°C in 20% glycerol) in R2A liquid media and used to inoculate 500 mL of NDW2 medium at 1% (v:v). After reaching mid-exponential phase (OD_600_ 0.3) aerobic growth at 30 ^o^C, the culture was pelleted by centrifugation at ~6000 x g for 10 min, washed 2x and resuspended in 10 mL fresh NDW2 medium without electron donor. 5 mL of this final resuspension was stored in 200 mM NaOH for measuring protein content and the remaining 5 mL was introduced to the electrochemical cell, which already contained 50 mL NDW2 medium (acetate as electron donor) with the WE poised at +522 mV vs. SHE, after the abiotic current stabilized to a constant baseline. Filter-sterilized N2 gas was used throughout to maintain anaerobic conditions, and the poised WE acted as the sole electron acceptor. Cell densities were determined by plate counts (for *Delftia)* or DAPI staining (for *Azonexus).* The cell measurements techniques differed because of slow growth of Azonexus on synthetic NDW2 media plates. To measure protein content, samples were digested in 200 mM NaOH at 100 °C for 90 min, with vigorous vortexing at 15 min intervals. Soluble protein in the extracts was measured with Pierce BCA Protein Assay Kit (Thermo Scientific, CA, USA) using bovine serum albumin (BSA) as a standard.

### MICROSCOPY

For scanning electron microscopy (SEM), electrode samples were fixed overnight in 2.5% glutaraldehyde (Grade I, specially purified for use as an electron microscopy fixative, Sigm-Aldrich). Samples were then subjected to an ethanol dehydration series (25%, 50%, 75%, 90%, and 100% v:v ethanol, for 15 min each treatment) and critical point drying (Autosamdri 815 critical point drier, Tousimis Inc., Rockville, MD, USA). The samples were then mounted on aluminum stubs, coated with Au (Sputter coater 108, Cressington Scientific), and imaged at 5 keV using a JEOL JSM 7001F low vacuum field emission SEM. Samples were also imaged using fluorescence microscopy on a Nikon Eclipse Ti-E inverted microscope. Glutaraldehyde fixed samples were stained with FM 4-64FX (Life Technologies) membrane stain (5 µg/mL) and imaged using the TRITC channel (Nikon filter set G-2E/C).

## RESULTS AND DISCUSSION

### SAMPLING SITE

The Death Valley Flow System (DVFS) consists of highly fractured mostly carbonate-rock aquifers that form a regional groundwater flow system covering hundreds of square km; extending from recharge zones associated with Central Nevada Uplands to large discharge springs in the Amargosa Valley and Furnace Creek area of Death Valley (Winograd and Pearson, 1976; Belcher et al., 2009; Belcher and Sweetkind, 2010). The source of water used for this study, Nevares Deep Well 2, is located in the Death Valley portion of the discharge zone of this system and intersects the Furnace Creek Fault Zone, which is inferred to represent a major conduit for fluid flow through the Funeral Mountains (Belcher and Sweetkind, 2010; Thomas et al., 2013). Given the natural artesian flow associated with this site, the mostly uncased nature of the hole, and the fact that it was drilled without drilling fluid (see methods), NDW2 can be regarded as a unique window into a deep continental microbial ecosystem.

### CURRENT RESPONSE AND BACTERIAL COMMUNITY ANALYSIS IN ELECTROCHEMICAL ENRICHMENT

Anodic currents began developing within 20 days of the start of the electrochemical enrichment (Figure 2), at all working electrode conditions (WE1: 272 mV, WE2: 372 mV, WE3: 472 mV, and WE4: 572 mV vs. SHE). Separate testing with sterile media under identical operating conditions (abiotic control) did not result in any anodic current, indicating that the electron transfer was mediated by the resident microorganisms. The observed anodic currents were correlated with a modest decrease of electron donor (acetate) concentration, from 10 mM to 7.5 mM as measured by HPLC over a one month period, consistent with the activity of a microbial community that couples the oxidation of acetate to anodic reduction. Acetate was chosen as the electron donor in recognition of its important role as an intermediate in anaerobic degradation of organic matter, and in light of previous observations that acetate-oxidizing microorganisms were prevalent on anodes inserted into sediments (Wardman et al., 2014). Taken collectively, the overall increase in potentiostatic current, reduction in acetate concentration and presence of cells on the working electrodes, point to the enrichment of a microbial community capable of electron transfer to the reactor’s working electrodes across a wide range of oxidizing potentials. This was also consistent with SEM images of the working electrodes, which revealed the formation of biofilms containing various cellular morphologies (Figure 3).

**Figure 2.**
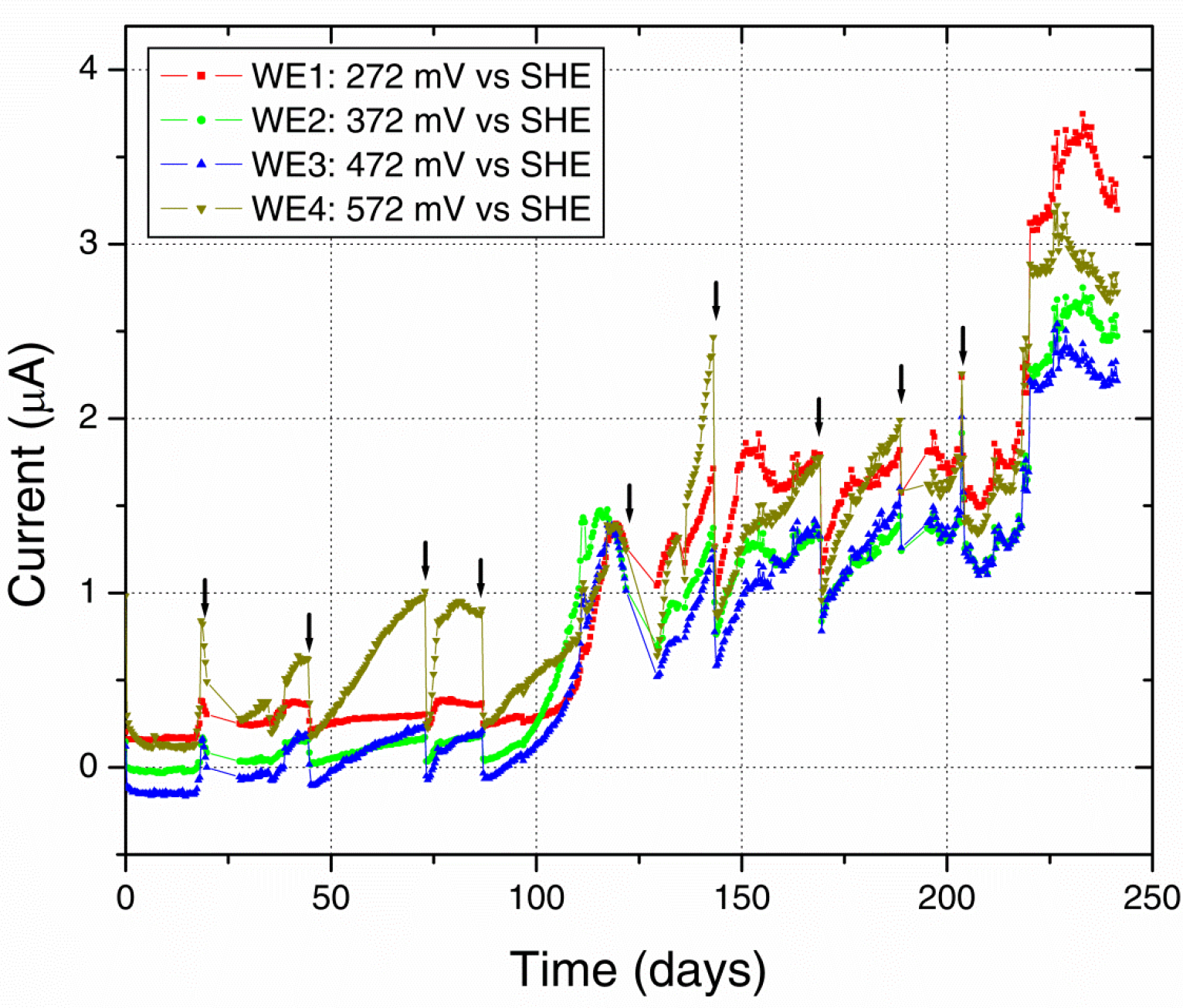
**Electrochemical enrichment** The increasing anodic current vs. time indicates the enrichment of microbial communities capable of mediating extracellular electron transfer to the working electrodes. Arrows indicate times of medium change, which resulted in an abrupt 3 decrease of anodic current before recovery of the increasing trend.

**Figure 3.**
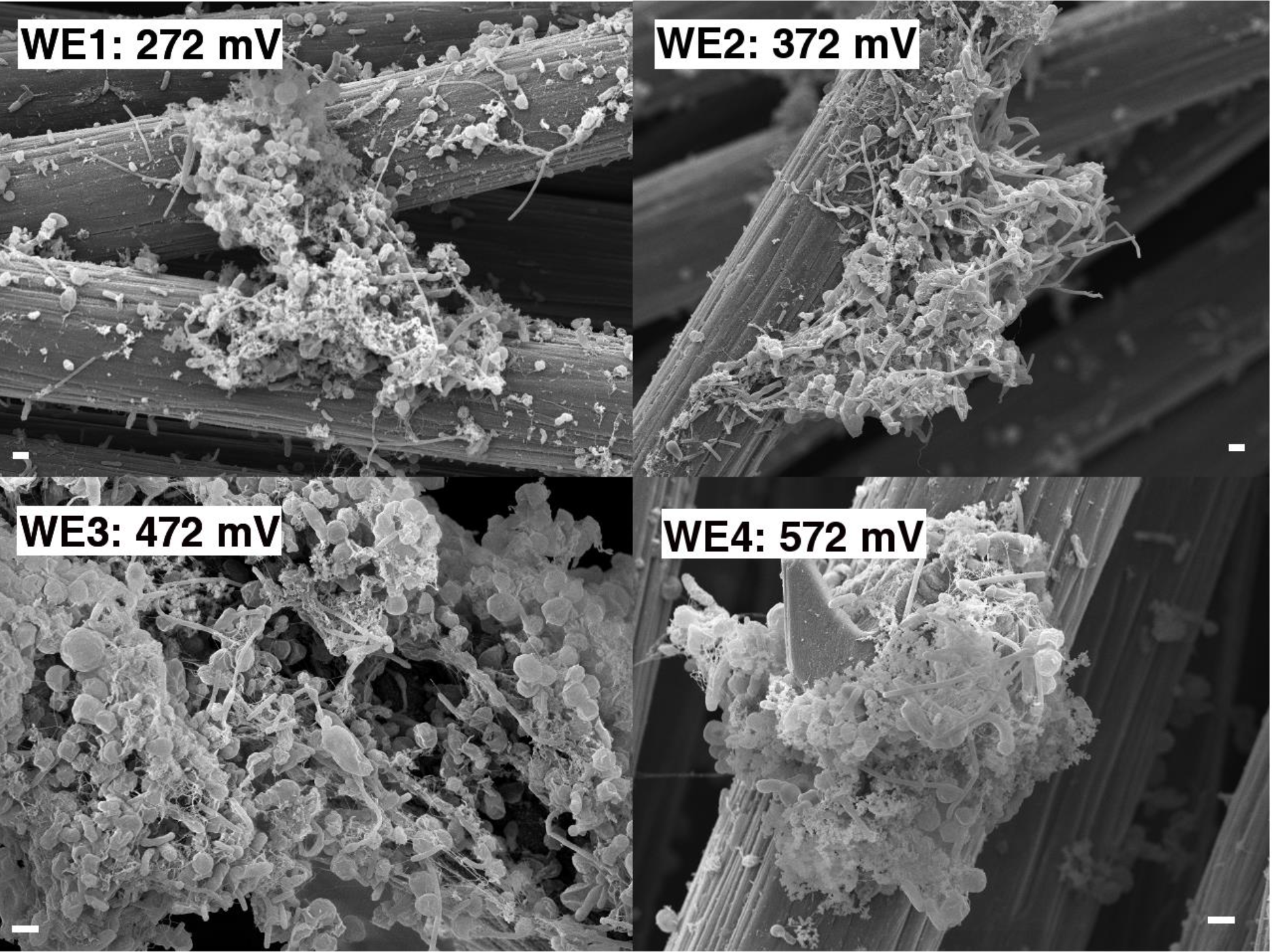
**Scanning electron microscopy of the enrichment electrodes and associated biomass (1 µm scale bar).**

Along with the increasing trend of anodic current over time, we observed an abrupt decrease in current with every change of medium (Figure 2). This may indicate a role for soluble components that act as redox shuttles carrying electrons from cells to the electrodes (indirect EET), as previously proposed for flavins secreted by *Shewanella* to mediate EET to electrodes (Marsili et al., 2008; von Canstein et al., 2008). While the observed decline in current with medium change is usually attributed to such redox shuttling mechanisms, it has alternatively been suggested that such a decline may originate from a decreased binding of the soluble redox components to cell-surface proteins that perform direct EET to external surfaces (Okamoto et al., 2013). While the precise mechanism(s) of cell-to-anode electron transfer in our enrichment culture is more difficult to pinpoint compared to the canonical metal-reducing model systems such as *Shewanella,* it is interesting to also note that SEM imaging (Figure 3) revealed the presence of extracellular filaments morphologically similar to bacterial nanowires (Gorby et al., 2006).

The microbial community analyses, performed using 16S rRNA gene pyrosequencing, revealed shifts between the initial samples, the DHS and batch enrichments used as inocula for the electrochemical reactor, and the communities attached to the electrodes. The original groundwater inoculum was dominated by members of the *Proteobacteria* (78%), with presence of *Nitrospirae* (10%), *Firmicutes* (2.5%), *Chloroflexi* (2.3%), *Spirochaetes* (1.24%), and other unclassified organisms (2.7%). The sequences obtained from the DHS effluent belonged primarily to *Alphaproteobacteria* (60%) and *Betaproteobacteria* (16%), in addition to *Actinobacteria*> (7%) and *Bacteriodetes* (5%). The Fe(III)-reducing batch culture used as inoculum for electrochemical reactor resulted in an increase of *Betaproteobacteria*, and this trend continued in the electrochemical enrichment with all electrodes dominated by the *Comamonadaceae* and *Rhodocyclaceae* families in the *Betaproteobacteria* (Figure 4). Clone libraries (approximately 40 clones) comprised of 21 clones from the genus *Delftia* (bootstrap 100%) and 1 clone from the genus *Acidovorax* (bootstrap 85%) within the *Comamonadaceae* family in addition to 8 clones from the genus *Dechloromonas* (bootstrap ranging from 68% to 95%) and 5 clones from the genus *Azonexus* (bootstrap ranging from 46% to 69%) within the *Rhodocyclaceae* family. One clone each was obtained from the genera *Ignavibacterium* (bootstrap 100%), *Mesorhizobium* (boostrap 100%) and *Aestuariimicrobium* (bootstrap 100%).

**Figure 4.**
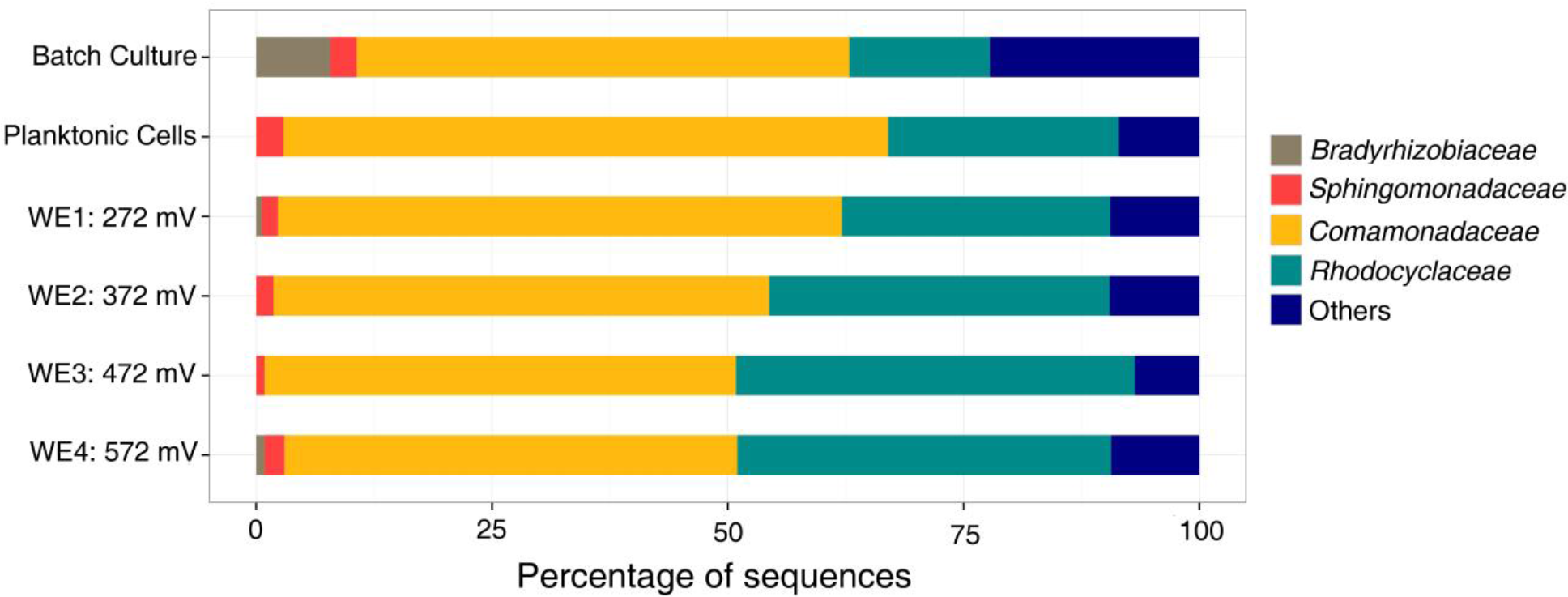
**Family level microbial community analysis of the electrochemical enrichment** Families were determined using 16S rRNA tagged pyrosequencing analysis of DNA extracts. A batch culture (acetate as electron donor and Fe(III)-NTA as electron acceptor) was the inoculum for the electrochemical reactor.

With acetate as the electron donor and electrodes poised within the redox potential range (300750 mV vs. SHE) of nitrogen-based redox couples like nitrate in neutral waters (Thauer et al., 1977), the electrochemical enrichment of *Comamonadaceae* and *Rhodocyclaceae,* reported here, is consistent with their previous description as short-chain fatty acid-utilising denitrifiers (Ginige et al., 2005; Timmers et al., 2012). *Rhodocyclaceae* are also known to reduce chlorate and iron (Hori et al., 2010; Sun et al., 2010). Indeed, both families have been observed on anodes in previous microbial fuel cell (MFC) enrichments (Timmers et al., 2012). Specifically, *Comamonadaceae* were previously found to dominate the anodic community of cellulose-fed MFCs inoculated with rumen microorganisms (Rismani-Yazdi et al., 2007), and *Rhodocyclaceae* dominated the anodic biofilm of an acetate-fed MFC enrichment inoculated with anaerobic digester slurry (Borole et al., 2009). In both cases, it was suggested that some members of these families could utilize electrodes as electron acceptors, in the absence of nitrate, while oxidizing short-chain fatty acids. However, no particular members of these families from these previous enrichments have been isolated or tested to confirm their EET ability. To date, the only confirmatory reports of EET from relevant pure cultures are limited to *Rhodoferax ferrireducens* (Chaudhuri and Lovley, 2003) and *Comamonas denitrificans* (Xing et al., 2010), from the *Comamonadaceae*, both of which have been shown to produce current at MFC anodes in the absence of alternate soluble electron acceptors (e.g. oxygen or nitrate). With this in mind, we proceeded to isolate representative members of the dominant groups in our electrochemical enrichment to directly test these pure cultures for electrochemical activity in half-cell reactors.

### ELECTROCHEMICAL MEASUREMENTS OF ISOLATED *DELFTIA* **and** *AZONEXUS* **STRAINS**

Two strains, designated WE1-13 and WE2-4, were isolated as pure colonies originating from the biomass associated with the 272 mV vs. SHE (WE1) and 372 mV vs. SHE (WE2) electrodes. Phylogenetic analyses of their 16S rRNA gene sequences demonstrated that these strains belonged to the genera *Delftia* and *Azonexus,* respectively. Remarkably, previous studies have hinted at potential EET activity from these genera, based on their minor appearance within MFC anode communities fed with glucose and acetate, respectively (Ishii et al., 2012; Chen et al., 2014). Since the mere presence of microbes in anode biofilms is not direct evidence of EET capability (Bond, 2010), these previous reports noted the need for additional studies targeting their electrochemical activity (Chen et al., 2014). Motivated by these reports, and the potential relevance of these microorganisms to subsurface environments by virtue of being enriched from source waters associated with a deeply-sourced spring, we tested *Delftia* sp. strain WE1-13 and *Azonexus* sp. strain WE2-4 as pure cultures in half-cell reactors. In addition to the reference and counter electrodes, each anoxic reactor contained a carbon cloth electrode (same material used in the electrochemical enrichment) poised at 572 mV vs. SHE, and acetate (10 mM) was used as electron donor.

Upon inoculation of *Delftia* sp. WE1-13 to reactors, the anodic current immediately started rising above the typical 50 nA background established from the sterile medium (Figure 5A), leveling off at 400 nA after 11 hr. This rise in current, indicative of microbial EET current to the anode, was accompanied by near doubling in total protein content (inset, Figure 5A) from 5.931 ± 0.109 mg (inoculum) to 10.645 ± 0.288 mg (sum of contributions from both planktonic cells and electrode-attached biomass after 12 hr). Similar chronoamperometry results were obtained from *Azonexus* sp. WE2-4, whose inoculation resulted in a slower rise of anodic current from a 70 nA sterile-medium background current to 325 nA after 60 hr (Figure 5B), concomitant with total protein increase from 5.487 ± 0.095 mg to 9.069 ± 0.301 mg (inset, Figure 5B).

**Figure 5.**
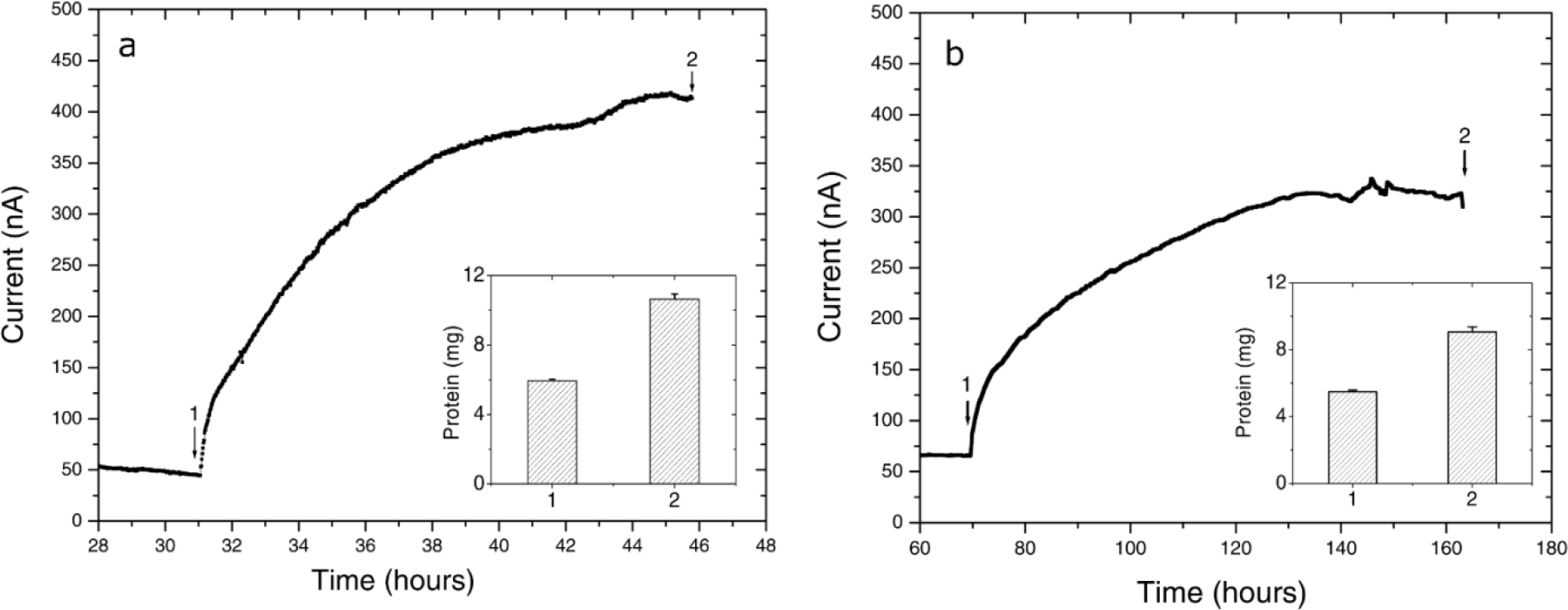
**Chronoamperometry of the isolated strains (A) *Delftia* sp. WE1-13, (B) *Azonexus* sp. WE2-4** In both cases, the working electrode was poised at 522 mV vs SHE, acetate (10 mM) was used as electron donor, and reactors were purged with N2 to maintain anaerobic conditions. Insets show the total protein content measured at time points labeled 1 and 2.

While different for the two strains, the time frame of the rise in anodic current is in both cases consistent with previous observations of electrochemically-active microbes (Chaudhuri and Lovley, 2003; Marsili et al., 2008; Fedorovich et al., 2009). However, the magnitude of anodic currents detected from the *Delftia* and *Azonexus* strains reported here is significantly lower than those typically reported for the metal-reducing microbes usually investigated as EET model systems. *Shewanella oneidensis* MR-1, for example, produced three orders of magnitude more current in reactors operated using identical electrode types and under similar conditions while using lactate as an electron donor (Xu et al., 2016). While this may be partially attributed to culturing and medium conditions that have not been systematically optimized for EET activity, the low currents observed here may point to the isolated strains’ lower ability to gain cellular energy from external redox active surfaces. The latter notion is consistent with our observation of very low cellular decay rates, but no growth, from the planktonic cell counts performed over the course of chronoamperometry. Specifically, the *Delftia* sp. WE1-13 cell density changed from 7 ± 2 x10^7^ CFUs/mL to 5 ±2 x10^7^ CFUs/mL, while *Azonexus* sp. WE2-4 showed a slight decrease from 6.9±1.4 x10^8^ CFUs/mL to 4.1±1.4 x10^8^ CFUs/mL (between the time points labeled 1 and 2 on Figure 5). Taken collectively, the small decrease in cell density, modest increase in protein content, and low EET currents observed, suggest low energy gain that may be directed for cell maintenance and decreased cellular decay, rather than growth. Considering the above, we heuristically compared EET rates the isolated strains relative to the well-characterized *S. oneidensis* MR-1 that is known to harness EET to gain energy for growth. Assuming an average bacterial protein content of 0.2 g/mL (Milo and Phillips, 2015) and average cell volume of *Delftia* and *Azonexus* as 0.4 µm^3^ (estimated from SEM images e.g. Figure 6) we calculated the average respiration rate of *Delftia* as 174.7 e^−^/s/cell (carbon cloth protein content ~1 mg; current ~350 nA) and of *Azonexus* as 326.4 e^−^/s/cell (carbon cloth protein content ~0.39 mg; current ~255 nA). These values are 3 orders of magnitude less than the respiration rate (6.2 x 10^5^ e^−^/sec/cell) observed for *S. oneidensis* MR-1 (Gross and El-Naggar, 2015).

We further investigated the impact of medium change on EET by *Delftia* sp. WE1-13, in light of the observation that the original enrichment’s anodic current decreased abruptly in response to this procedure (Figure 2). A separate reactor was inoculated with *Delftia* sp. WE1-13 after the establishment of a background sterile-medium current, as described above (Figure 6). Following the expected rise in current (up to 500 nA), during which the total planktonic cell density remained nearly constant (from 5±3 x10 to 6±1 x10 CFUs/mL), the spent medium and associated planktonic cells were replaced with fresh medium. Consistent with the enrichment’s observations (Figure 2), this resulted in an abrupt decrease of anodic current down to 60 nA followed by recovery to 200 nA over 6 hr (Figure 6A). At the conclusion of the experiment, the final total protein content (both planktonic and electrode biomass) was measured to be 12.764±0.466 mg, compared to the inoculum’s 6.128±0.115 mg. SEM and fluorescence imaging (Figures 6A and 6B) confirmed cellular attachment and the formation of a *Delftia* biofilm on the working electrode, consistent with measured protein from the electrode biomass. The drop in current upon medium change is consistent with loss of putative secreted components that may enhance EET by *Delftia* sp. WE1-13 to anodes. Alternatively, the medium change may have temporarily disrupted biofilm components that directly mediate electron transfer to the electrodes. These observations motivate further investigations into the identity of the charge carriers and precise pathway(s) involved. In this context, it is also interesting to note that the phylogenetically closely related *Delftia acidovorans,* an organism dominant in subsurface gold-associated mine communities (Reith et al., 2010), has been shown to produce secondary metabolites responsible for extracellular gold reduction and precipitation, providing protection from toxic soluble gold (Johnston et al., 2013). To check whether *D. acidovorans* is also electrochemically active, we obtained this strain from the German Resource Centre for Biological Material (DSMZ, DSM no. *39*), and tested it under identical conditions as *Delftia* sp. WE1-13. Our tests revealed that *D. acidovorans* is indeed electrochemically active (data not shown) with similar anodic currents, cell survival, and modest increase in protein content. Clearly, additional studies are required to pinpoint both the precise mechanism of EET, and its relevance to cellular protection and energy generation in subsurface environments.

**Figure 6.**
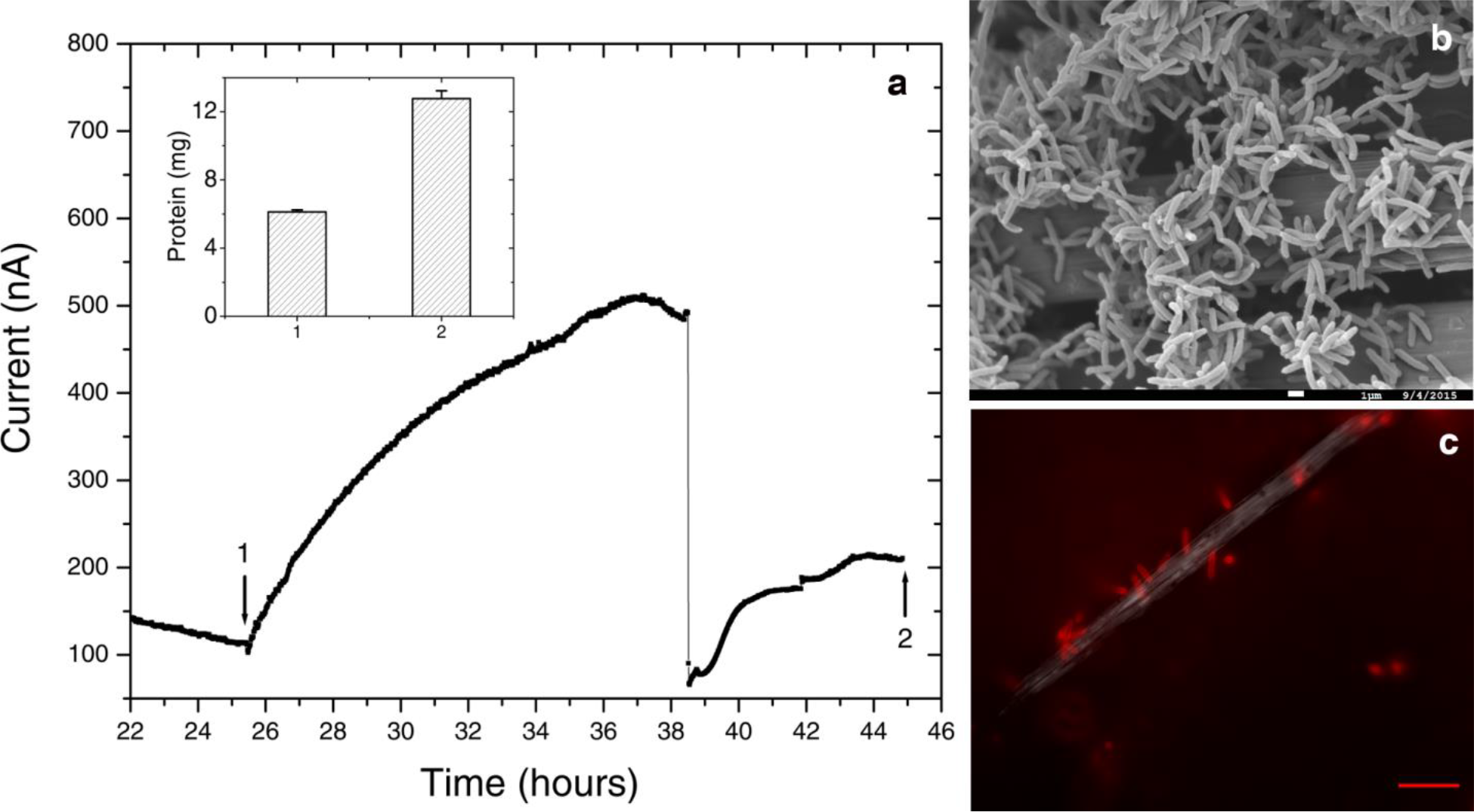
**Further electrochemical and microscopic characterization of *Delftia* sp. WE1-13** (A) Medium change resulted in a sudden decrease of anodic current, as observed in the enrichment cultures. Inset shows total protein content measured at the time points marked 1 and 2. B) Scanning electron microscopy (1 µm scale bar) shows attachment of the rod-shaped of *Delftia* sp. WE1-13 on the carbon cloth fibers. (C) *In vivo* fluorescent image (FM 4-64 FX membrane stain) confirms the attachment of *Delftia* sp. WE1-13 cells to the electrode during electrochemical analysis (10 µm scale bar).

### IMPLICATIONS

While alternative anaerobic metabolisms (e.g. nitrate reduction) by *Delftia* and *Azonexus* species were previously known, our study provides direct evidence that these organisms can also gain some energy for survival and/or persistence by electron transfer to external surfaces. The electrochemical enrichment and isolation of these organisms from a fractured rock aquifer raises interesting questions about the relevance of EET in energy-limited subsurface environments, where the diverse host rocks may provide insoluble electron acceptors (or electron donors) that greatly expand the range of redox couples available for the resident microorganisms. Significantly, these modes of energy acquisition from external substrates, detectable using electrochemical enrichment and analytical techniques, are easily missed using traditional cultivation strategies. In addition to helping identify new microbial candidates for renewable energy and biocatalytic applications, *ex situ* and *in situ* electrochemical approaches expand the range of cultivation conditions available for subsurface microorganisms, and may shed light on potential mechanisms for cellular survival and detoxification activity in these environments. It should be noted that while the focus here was on a continental subsurface site, similar relevance of EET and electrochemically-active microorganisms may be of interest in the vast marine subsurface or even in environments on extraterrestrial planetary bodies.

## ACKNOWLEDGMENTS

The authors thank Dr. Lewis Hsu for input on the electrochemical enrichment reactor design, Dr. Brandi Kiel Reese for guidance on sequencing and analysis, and Dr. Shuai Xu for helpful comments on electrochemical measurements of the isolates. The authors are also grateful for Dr. Annie Rowe’s comments on the manuscript. Thanks also to Michael King and others from the Hydrodynamics Group LLC, for information concerning the drilling and completion of Nevares Deep Well 2 and to Richard Friese, Genne Nelson and others from the National Park Service for permits and assistance with sample collection. Samples were collected under special use permit DEVA-2009-SCI-0005, issued to DPM. Scanning Electron Microscopy was performed at the University of Southern California’s Center of Excellence in Electron Microscopy and Microanalysis. YJ is grateful for a 2015-2016 merit fellowship from the USC Women in Science and Engineering (WiSE) program. This work was supported by the NASA Astrobiology Institute (Life Underground project) under cooperative agreement NNA13AA92A.

